# The serotonin 1B receptor modulates striatal activity differentially based on behavioral context

**DOI:** 10.1101/2025.10.21.683729

**Authors:** Ka H. Ng, Arati Sharma, Katherine M. Nautiyal

**Affiliations:** Department of Psychological and Brain Sciences, Dartmouth College, 6207 Moore Hall, Hanover, NH, 03755 USA

## Abstract

The dorsomedial striatum (DMS) is critical for goal-directed behavior and has been implicated in both motivating and inhibiting behavioral responses. The DMS circuitry is complex as it integrates multiple inputs from the cortex, thalamus, and other subcortical structures including midbrain dopamine neurons. Though less studied, serotonin neurons from the dorsal raphe nucleus also richly innervate the DMS, which expresses the majority of the 14 receptor subtypes for serotonin. In particular, slice electrophysiology shows that the serotonin 1B receptor (5-HT1BR) impacts DMS physiology and plasticity, and behavioral experiments show that 5-HT1BR expression modulates impulsivity and other DMS-dependent reward-related behaviors. In these studies, our goal was to investigate the effects of 5-HT1BR on the DMS in vivo during goal-directed behavior in mice. Using a genetic 5-HT1BR loss-of-function model, we examined the calcium activity of individual medium spiny neurons (MSNs) in the DMS during operant tasks of responding and waiting. We found that knockout of 5-HT1BRs resulted in a significant reduction of excitatory calcium responses and an increase in the proportion of cells with inhibitory calcium responses following receipt of a reward. This suggests that serotonin may recruit MSN activity in response to reward via actions at 5-HT1BRs. On the other hand, in a behavioral paradigm designed to test impulsivity, we found that serotonin may inhibit DMS calcium activity through 5-HT1BRs. Specifically, mice lacking 5-HT1BR expression had a larger proportion of cells showing increased calcium responses during the waiting period of the trial, compared to controls. These results point to the importance of in vivo studies to understand the functional role of DMS serotonin in reward-related behavior. Overall our results demonstrate that serotonin can modulate the DMS in a behavioral state-specific manner, potentially providing a mechanism for how serotonin effects on behavior are context-dependent.

## Introduction

The dorsomedial striatum (DMS) is critical for learning and executing goal-directed behavior across species. Its role in action-outcome learning has been extensively studied, and research has demonstrated the importance of dopamine modulation of DMS control of reward-motivated behavior (Dayan & Balleine, 2002). However, the DMS is also richly innervated by serotonin afferents from the dorsal raphe nucleus (DRN), and the majority of the 14 serotonin receptor subtypes are expressed in this region (Di Matteo et al., 2008; Nair et al., 2020a; Soghomonian et al., 1989). Additionally, evidence shows that serotonin is critical for goal-directed decision making, as disruption of serotonin signaling in humans and mice impairs inhibition of responses to devalued rewards, and shifts behavior toward more habitual responding (Ohmura et al., 2021; Worbe et al., 2015). This suggests that serotonin may be involved in recruiting DMS-related circuitry to engage in model-based decision making. However, limited research has investigated how serotonin modulates DMS circuitry in vivo.

Many convergent lines of research support a role for serotonin in the modulation of DMS signaling (Mathur & Lovinger, 2012; Nair et al., 2020b). Using a GRAB-5-HT biosensor in vivo, we showed that serotonin encodes an anticipatory value-graded reward signal in the DMS (Spring & Nautiyal, 2024). Additionally, serotonin can modulate the release of dopamine in the DMS (Sershen et al., 2000). More directly, electrophysiology studies show that serotonin increases excitation of cholinergic and fast spiking interneurons in the DMS, and reduces local collateral inhibition between medium spiny neurons (MSNs; Blomeley & Bracci, 2009; Pommer et al., 2021; Virk et al., 2016). Finally, serotonin has the potential to influence plasticity in the DMS by inducing long-term depression of cortical and thalamic inputs to MSNs (Cavaccini et al., 2018; Mathur et al., 2011). Interestingly, the majority of these effects of serotonin on DMS physiology have been shown to be mediated through signaling at the serotonin 1B receptor (5-HT1BR).

In addition to the evidence supporting a role for 5-HT1BR modulation of DMS physiology, many studies show that 5-HT1BRs also influence DMS-dependent behavior. Whole brain 5-HT1BR knockout (KO) mice display increased incentive motivation, impulsive, and compulsive behavior (Desrochers et al., 2021; Nautiyal et al., 2015; Rocha et al., 1998). However, 5-HT1BRs are expressed on axon terminals widely throughout the brain, and the region and cellular specificity for the effects of 5-HT1BR on these behavioral phenotypes are still being worked out. Tissue-specific knockout of 5-HT1BRs in the DMS has been associated with altered reward-seeking phenotypes (Li et al., 2021; Virk et al., 2016). Though interestingly this may arise through a number of different mechanisms via 5-HT1BR expression on a variety of distinct cell types within the DMS. These include heteroreceptor-mediated inhibition of acetylcholine release from striatal cholinergic interneurons, glutamate release from cortical inputs, and MSN collaterals inhibition within the DMS (Burke & Alvarez, 2022; Li et al., 2021; Pommer et al., 2021; Virk et al., 2016). Additionally, 5-HT1B autoreceptors are located on presynaptic terminals of serotonin inputs to the DMS allowing serotonin to inhibit its own subsequent release. The heterogenous localization of 5-HT1BR expression within the complex DMS local circuitry highlights the difficulty of understanding serotonin effects on DMS activity, even just through one of its 14 receptors.

In this study, we set out to understand how global deletion of 5-HT1BR expression influences the calcium activity of DMS neurons during action and inhibition. We were specifically interested in understanding if the encoding of action or waiting in the DMS were altered in the absence of 5-HT1BRs. We used a one photon head-mounted miniature microendoscope to image the calcium activity of MSNs in the DMS neurons in mice during two operant tasks to examine action and inhibition. The first was a time-pressured cue-guided response task in which mice had a limited cue period to respond for a reward. In this task, we found that mice lacking 5-HT1BR expression had less excitatory calcium responses suggesting that serotonin may increase MSN reward-related neural activity via 5-HT1BRs. In a second task, mice were challenged to wait during a delay period, before responding for a reward. Interestingly, during the delay period of this task, mice lacking 5-HT1BRs showed increased recruitment of excitatory calcium activity compared to controls suggesting that 5-HT1BR activation could inhibit MSN activity during behavioral inhibition. The differential effects of 5-HT1BR expression on MSN calcium activity during action and waiting suggest that serotonin can have different physiological effects on a population of MSNs depending on behavioral context. These results establish possible mechanisms of 5-HT1BR-dependent modulation of DMS-dependent reward-related behavioral dysfunction and point to the importance of studying the effect of serotonin in the brain in vivo.

## Methods

### Animals

All mice used for these experiments were bred in the Center for Comparative Medical Research at Dartmouth College. Male and female mice lacking expression of 5-HT1BR (N=9) and littermate genetic controls (N=8) were bred by crossing tetO1B+/+::βActin-tTS+/-males to tetO1B+/+::βActin-tTS-/-females as previously reported (Nautiyal et al., 2015). Mice were weaned at postnatal day (PN) 21 into cages of 2–4 same sex littermates and maintained on a 12:12 light-dark cycle on *ad libitum* chow and water until experimental testing began as described. All procedures were approved by the Dartmouth College Institutional Animal Care and Use Committee.

### Surgical procedures

Mice were anesthetized with isoflurane (1-3%) in 0.8ml/min oxygen, and headfixed into a stereotax (Kopf). The scalp was cleaned, shaved, and opened, and the connective tissue was cleared from the skull. The skull was leveled and a 1.0mm burr hole was drilled at bregma (AP+0.8; ML+1.6mm). The dura was removed, and the cortical tissue under the craniotomy was aspirated. AAV9.CamKII-GCaMP6f.WPRE.SV40 (2.5 x10^12^ GC/ml) was injected into the parenchyma in two separate infusions of 250nl each at a rate of 50nl/min at 3.0 and 2.6mm ventral to dura using a glass microsyringe (Hamilton) driven by a programmable pump (World Precision Instruments). A gradient refractive index (GRIN) lens (1mm diameter, 4mm long, Inscopix) was lowered into the brain to a depth of 2.3mm ventral to dura and affixed to the skull with skull screws and Metabond (Parkell). Mice were individually housed following surgery. Approximately 6-8 weeks later GCaMP fluorescence was visualized with an nVista miniature microscope (Inscopix) and a baseplate was cemented to the headcap with additional Metabond cement.

### Behavioral training and testing

The DIY-NAMIC system (Lee et al., 2020) was used for behavioral training and testing. Mice were initially habituated to retrieve water in their homecage from the center noseport. A solenoid delivered 10ul of water through a blunted 18 gauge needle upon each head entry. During the first 4 days, a reward retrieval training paradigm (P1) was presented in which the cue light in the center port was illuminated and mice received 10 μl of water upon nosepoking into the center port. After water retrieval, a variable intertrial interval (ITI), with an average of 45 s preceded the next light illumination. Next, mice were trained for 3-4 days on a second paradigm (P2) which required a nosepoke to either side port when illuminated with a cue light. Mice were then trained for 4 days on a third paradigm (P3) to self-initiate trials through nosepoke responses to a blinking light in the center port which indicated trial availability after the ITI. This illuminated both left and right cue lights. A nosepoke response to either side port was rewarded. Mice were then trained for 3 days on a fourth training paradigm (P4) in which only one of the two side ports was illuminated following trial initiation, and a nosepoke only to that port resulted in reward delivery. Incorrect nosepokes to the unlit port terminated the trial without reward and started the ITI. Subsequently, the cue duration was shortened to 5s (P5), and then 1.5s (P5b) for 2 days each. One mouse did not reach criteria for behavioral performance during this time-pressured 1.5s cue and was excluded from analysis for this paradigm. Mice were imaged in the P5b paradigm for 1-2 days to measure the calcium activity of MSNs in the DMS during action and reward consumption. Subsequently, a delay of 3s was introduced between trial initiation and the cue response in order to examine the calcium activity during waiting behavior. Mice were run on this paradigm (P6.3) for six days, with imaging on the first and sixth session. Two mice did not reach criteria for behavioral performance and were therefore excluded from further testing and analysis of calcium imaging data. Throughout training and testing, all port entries were detected with infrared sensors and sent through the Arduino UNO to Processing software on a linked PC. All behavioral programs (P1-P6.3) and processing scripts are available online at https://github.com/DIY-NAMICsystem. Following these training paradigms, the ports were blocked off in their homecage to restrict access to allow for imaging during behavior within a discrete session.

### Data Acquisition

Imaging was performed in 40 min long sessions by affixing the one photon Inscopix nVista miniature microscope to the baseplate and removing the access barrier to the ports. Data was recorded using the nVista acquisition software at 20 frames per second with 50ms exposure. Optimal LED settings (0.5-0.6 mW/mm^2^ range) were selected for each mouse and remained consistent for each mouse over all imaging sessions. The Arduino UNO that controlled the DIY-NAMIC system initiated the calcium imaging session by sending a 5V start signal to the Inscopix Data Acquisition Software (IDAS). The Arduino UNO also emitted trains of 5s on/off 5V signals collected by an IDAS GPIO port. These pulses also allowed synchronization of the overhead camera video recording used for motion tracking. Additional 5V signals were also sent to the IDAS GPIO for all DIY-NAMIC infrared port entries to validate signal alignment. Video footage was captured using an overhead camera (ELP-SUSB1080P01-LC1100) at 30 frames/second.

### Calcium Imaging Processing

Raw videos were spatially cropped to remove areas not in the field of view of the lens, and spatially downsampled by a factor of 3. The videos were then processed using the python implementation of the Calcium Imaging Analysis package, CaImAn (Giovannucci et al., 2019). Motion correction was performed using the NoRmCorre algorithm (Pnevmatikakis & Giovannucci, 2017). Parameters for motion correction for each video were selected using a grid search approach, where multiple iterations of the model were tested using different combinations of parameter values, and optimized for stability of the video frames over the session. Spatial contours and temporal fluorescence traces for putative cells were identified using the constrained non-negative matrix factorization (CNMF-e) algorithm (Zhou et al., 2018). Parameters for CNMF-e were selected for each video using a grid search approach. The optimal set of parameters were selected based on: the dimensions of spatial footprints, their spatial location in the field of view, the correlation of spatial footprints within a session, and the signal to noise ratio of the temporal traces. The components obtained using the optimal combination of parameters were then sorted manually by two observers to remove components with signal dynamics or spatial characteristics that are inconsistent with neuronal calcium activity. The data from one mouse in a P5b imaging sessions yielded fewer than 10 cells and was therefore excluded from analysis for that paradigm. Delta F/F videos were generated using a minimum z-projection image of the video. Calcium transients were extracted from the denoised calcium fluorescence traces using the sparse non-negative deconvolution algorithm (OASIS) implemented in CaImAn (Friedrich et al., 2017).

### Behavioral Data Processing

The arduino UNO behavioral data was aligned to the Inscopix GPIO data using the 5s on/off pulse to ensure continued alignment throughout the session. Pixel changes per frame were quantified using ezTrack (Pennington et al., 2019) and then thresholded at a 10 gray scale unit to remove baseline pixel fluctuations. The locomotion signal was then binned into 20 frames per seconds and aligned to the neural and Arduino data. For the action task, locomotion was compared between genotypes over the whole trial using a 2-way ANOVA (genotype by time). For the waiting task, locomotion data was compared between genotypes and session across different phases of the trial using mixed effect models using the lmer function in RStats.

### Analysis

For each cell, the last 6s of every ITI (prior to trial availability) was averaged establish average baseline activity. Cells were assessed for significant changes in calcium event rate during four time epochs of correct trials relative to the baseline period by 10,000 round paired permutation (trial initiation, cue presentation, reward presentation, post-reward period). Correction for multiple comparisons were performed using Bonferroni to account for multiple trial epochs. Average calcium event traces were calculated by averaging the traces for each cell across all animals within each genotype and then compared using a 2 way repeated measures ANOVA (genotype x time). Percentages of inhibited, excited, and non-responsive cells were compared between genotypes using a Chi square test for each trial epoch.

### Lens placement

Following all behavioral testing, mice were anesthetized with Ket/Xyl and transcardially perfused with 0.9% PBS followed by 4% paraformaldehyde (PFA). Skulls were removed and postfixed overnight in 4% PFA, before the brain was removed and then postfixed again in 4% PFA overnight. Brains were then cryoprotected in 30% sucrose in PBS, and then frozen for sectioning at 40 um on a cryostat (Leica CM3050S). Sections were stored in PBS and immunostained with chicken anti GFP (ab13970, lot GR3190550-15, Abcam) followed by cy2 anti-chicken (Jackson ImmunoResearh Laboratories, Cat# 703-546-155) and DAPI (Thermo Scientific, Cat# 62248). Stained sections were mounted onto slides and coverslipped with Krystalon for verification of lens placement and GCaMP6f expression (Fig 1A). Localization of the lens for all mice with reported data was plotted on the brain atlas sections corresponding to 1.34 and 0.74 mm rostral to bregma (Fig 1B; Franklin & Paxinos, 2007).

**Figure 1.**
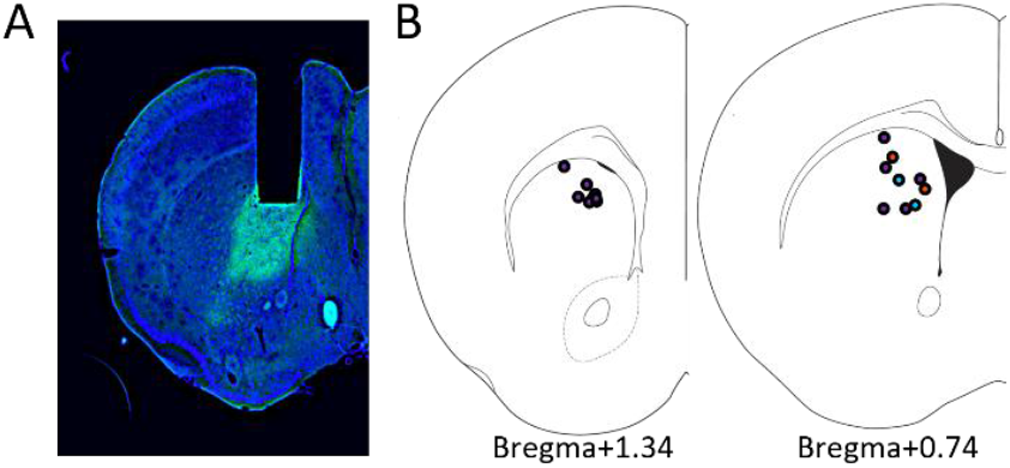
Localization of calcium imaging in the DMS. (A) Representative photomicrograph showing the location of the GRIN lens track and GCaMP6f expression (green) on a DAPI stained (blue) coronal section. (B) Atlas sections show reconstructed localization of the lens placement from each imaged mouse. Blue indicates mice excluded from analysis in the action task, and orange indicates mice excluded from analysis for the waiting task (see methods for details).

## Results

All mice (N=6 Controls, N=9 5-HT1BR KOs) were trained in a cue-guided operant response task (Fig 2A). Mice performed an average of 45.9±0.7 trials per session with 59±3% of the trials rewarded, 5±1% incorrect, and 37±3% percent omitted (Fig 2B). The high omission rates are a reflection of the time pressure in this task design which allows a cued response time period of only 1.5s following trial initiation. There were no significant effects of 5-HT1BR genotype on any behavioral measures in this task, including the total number of trials initiated (t_13_=1.51, p=0.155), percentage of correct trials (t_13_=1.62, p=0.128), or latency to respond to the cue (avg 0.93±0.3s; t_13_=1.59, p=0.136). There were also no significant differences in body weight (t_13_=1.46, p=0.169) or overall locomotor activity during the trial (F_1, 13_=1.90, p=0.191) between the genotypes. Calcium activity was recorded in 537 and 842 cells in control and 5-HT1BRKO mice, respectively. Activity for each cell was averaged over all rewarded trials within a session and aligned to the operant nose poke response. Cellular calcium activity heatmaps show that this task recruits the DMS in both genotypes, as seen by enriched calcium activity around the operant response in the correct trials compared to a baseline period during the ITI (Fig 2C).

**Figure 2.**
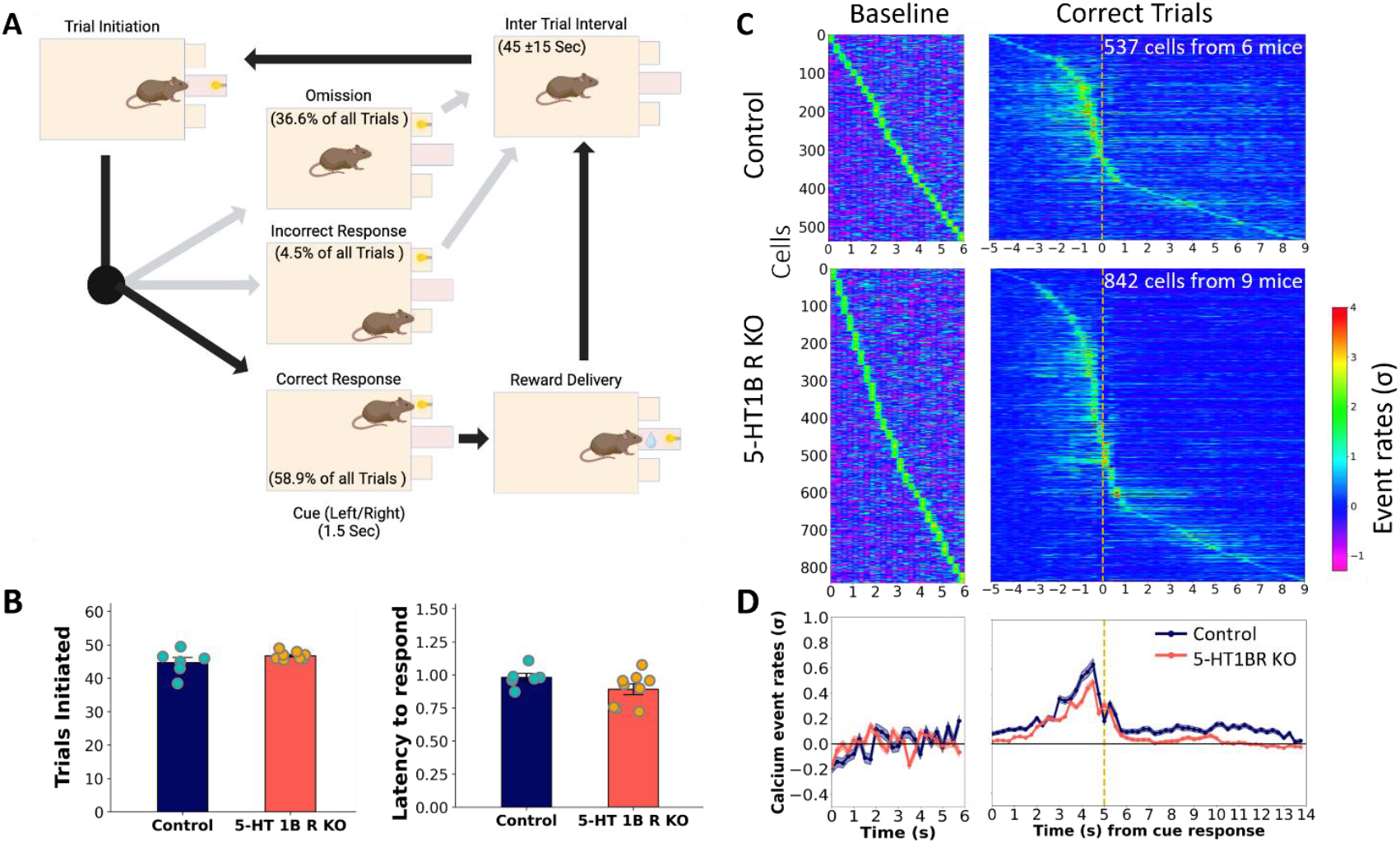
DMS calcium activity during an appetitive instrumental task. (A) Schematic showing operant trial structure including average percentage of trial types (omission, incorrect and correct responses) across all mice. (B) Behavioral data showing the average number of trials initiated (left) and average latency to respond to the cue on correct trials (right) for controls and mice lacking 5-HTlBRs. (C) Normalized calcium event rates for cells averaged over all trials during ITI baseline (left) and correct trial (right) for Control (top) and 5-HT1BR KO mice (bottom), sorted by time of peak event rate. (D) Normalized event rates are shown averaged for all cells over baseline and trial periods for Control (blue) and 5-HT1BR KO (orange) mice.

### 5-HT1BR promotes reward-related increases in neural activity in the DMS

During the ITI baseline, there were no effects of 5-HT1BR expression on calcium event rates (Fig 2D, t_1377_= 0.83, p = 0.41). During the trial, although population traces revealed qualitatively similar patterns of calcium activity between genotypes, there were notable differences in the magnitude of the average calcium event rates. Collapsed across the entire trial, there were significantly higher calcium event rates in cells from control mice compared to those from 5-HT1BR KO mice (t_1377_=7.96, p<0.0001). To further investigate the source of the decreased calcium event rates in the population-averaged traces in mice lacking 5-HT1BR across the extent of the trial, individual cells were characterized as excitatory or inhibitory in each of four epochs (trial initiation, cue response, reward consumption, and post reward consumption) of the task compared to the ITI baseline activity. This revealed significant effects of 5-HT1BR expression on the balance of excitation and inhibition of DMS calcium activity during this task (Fig 3A). Specifically, the percentage of cells that showed excitatory responses during the trial was larger in controls compared to 5-HT1BR KO mice in all four epochs of the trial (χ^2^(1, N=930)>6.07, p<0.013). Additionally, there was a higher percentage of cells showing significant decreases in calcium event rates in 5-HT1BR KO mice during the reward and post reward periods (χ^2^(1, N=930)>5.62, p<0.018). These data suggest that the absence of 5-HT1BR expression results in decreased activity in MSNs in the DMS during goal-directed behavior via less excitation and more inhibition.

**Figure 3.**
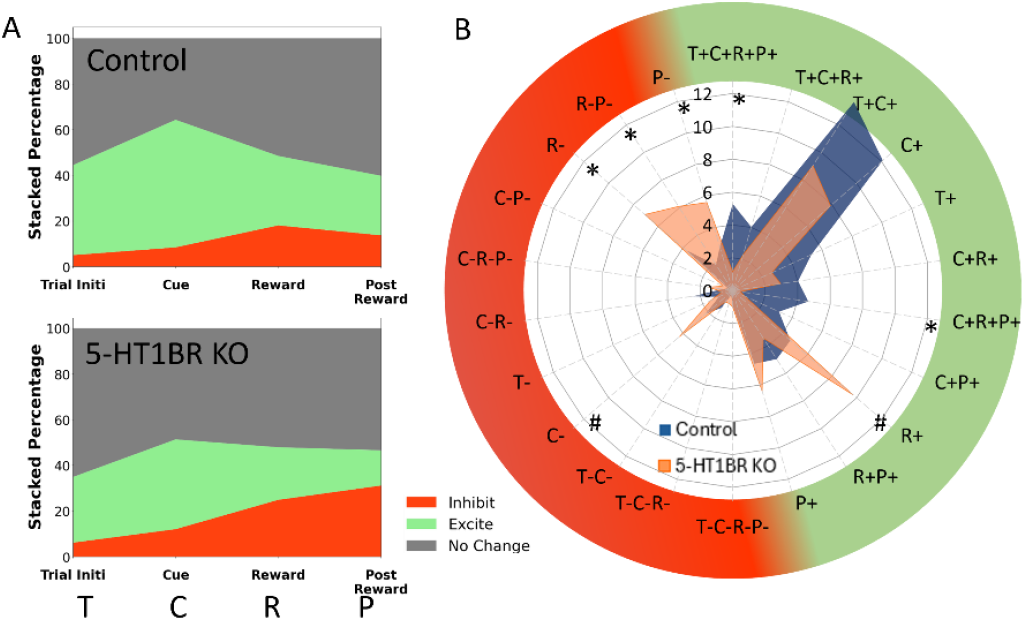
5-HT1BR influences the balance of excitation and inhibition during goal directed behavior. (A) The percentage of cells showing no change (grey), increased {green), or decreased (red) calcium activity during each of four trial epochs (trial initiation, cue presentation, reward presentation, post reward consumption) compared to baseline are shown for control (top) and 5-HT1BR KO (bottom). (B) Radial plot shows the percentage of responsive cells according to their mixed selectivity across all four trial epochs for control (orange) and 5-HT1BR KO (blue) mice. *, p<0.05; #, p<0.07

Since these initial cell categorizations did not consider mixed selectivity over different trial epochs, we further classified cells according to combinations of trial epochs that cells showed significant excitation or inhibition of calcium event rate above baseline ITI levels. Twenty-two categories accounted for over 80% of responsive cells (Fig 3B), and interestingly, the percentage of cells across categories were different between genotypes (category x genotype interaction: F_21,546_=1.62, p=0.039). Specifically, control mice showed a significantly higher percentage of cells excited during the trial (T+C+R+P+: p=0.014 and C+R+P+: p=0.020), while 5-HT1BR KO mice showed a higher percentage of cells inhibited during the reward epochs of the trial (R-: p=0.028, R-P-: 0.017, and P-: p=0.035). This suggests that in the absence of 5-HT1BR expression, the decreased excitation may be task-related, while the increased inhibition is relatively later in the trial, after the operant response, and potentially specific to the reward-related responses.

When examining reward-related calcium activity, regardless of mixed-selectivity, there was a higher proportion of neurons that were reward-responsive (either excited or inhibited) in mice lacking 5-HT1BR compared to controls (Fig 4A; 39% in controls and 53% in 5-HT1BR KOs; χ^2^(1, N=1379)=27.99, p<0.0001). Expression of 5-HT1BR significantly influenced the excitation-inhibition balance in trial epochs following the operant response to the cue (Fig 4B; F_5, 156_=6.07, p<0.0001 for genotype x cell type interaction). This was most notable in the number of cells showing significant excitation during reward receipt and continuing in the post-reward period, which was reduced in mice lacking 5-HT1BRs (R+P+; p=0.005). Not only did a lower percentage of cells show excitation in 5-HT1BR KO mice, but population calcium traces show that cells in 5-HT1BRKO mice also had lower calcium event rates on average during the reward and post-post reward epochs (Fig 4C; R+: p=0.0014, P+: p=0.013; R+P+: p=0.0421). In addition to the decreased excitation, there was also increased inhibition seen following reward consumption in terms of a higher percentage of cells showing inhibitory responses during the post-reward period in 5-HT1BR KO mice (Fig 4B; P-; p=0.008). This was not associated with any differences in average amplitude of the inhibition during the reward or post reward epochs (Fig 4C; P-: p=0.136). Overall, the calcium data from this instrumental task suggests that 5-HT1BR may mediate a reward-related increase in neural activity in the DMS, which is consistent with our previous data showing that 5-HT1BR expression influences reward-sensitivity phenotypes (Desrochers et al., 2021).

**Figure 4.**
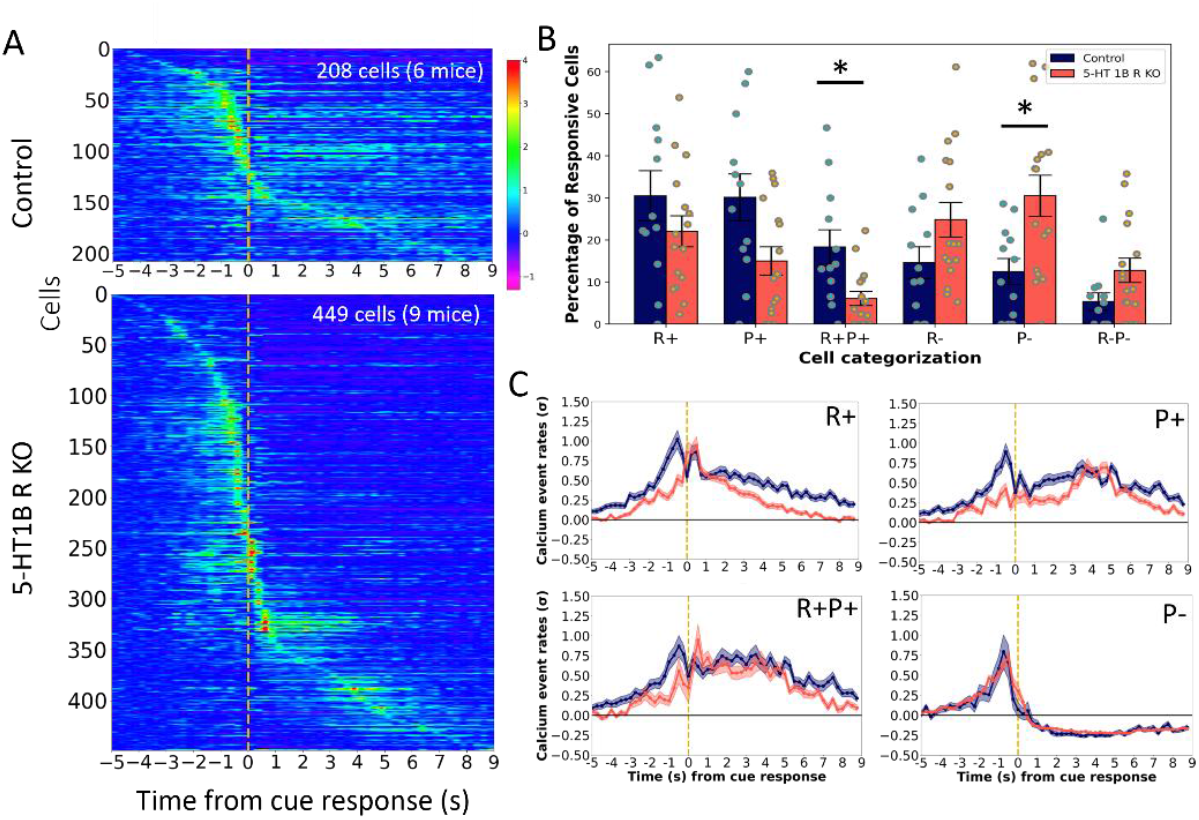
5-HT1BR expression influences DMS activity associated with reward. (A) Normalized calcium event rates are plotted for cells showing a significant response to reward for control (top) and 5-HT1BR KO (bottom) mice relative to cue response (yellow dashed line). (B) The percentage of responsive cells that are responsive to the reward and post reward epochs, regardless of their responses to trial initiation or cue presentation, are shown for both genotypes. (C) The average calcium activity is shown for cells classified as excited during the reward period (R+), post reward period (P+), or both (R+P+), or inhibited during the post reward period (P-) for control (blue) and 5-HT1BR KO (orange) mice. *, p<0.05

### 5-HT1BR inhibits waiting-related increases in neural activity in the DMS

We used a second operant paradigm to probe the role of serotonin in the DMS during waiting by inserting a 3s delay following trial initiation before the cue light was illuminated for responding (Fig 5A). Given that this paradigm was challenging for the mice, mice were rewarded on an average of only 14±3% of the trials they initiated on their first session. However, there were no genotype differences in the number of trials initiated (F_1,13_=0.14, p=0.712), or percentage of rewarded trials (F_1,13_=0.44, p=0.520). Additionally, consistent with our previous data showing increased impulsivity in mice lacking 5-HT1BR (Desrochers et al., 2021; Nautiyal et al., 2015), performance in this task, as measured by the percentage of trials with premature responses, was worse in 5-HT1BR KOs compared to controls (main effect of genotype F_1,13_=6.84, p=0.021). 334 and 645 cells were analyzed from control and 5-HT1BR KO mice while they performed this waiting task (Fig 5C). There were no significant genotype differences in the number of cells identified per mouse (F_1, 13_=0.36, p=0.561), nor in the calcium events for all cells over the trial (F_1, 977_=0.08, p=0.772; Fig 5D).

**Figure 5.**
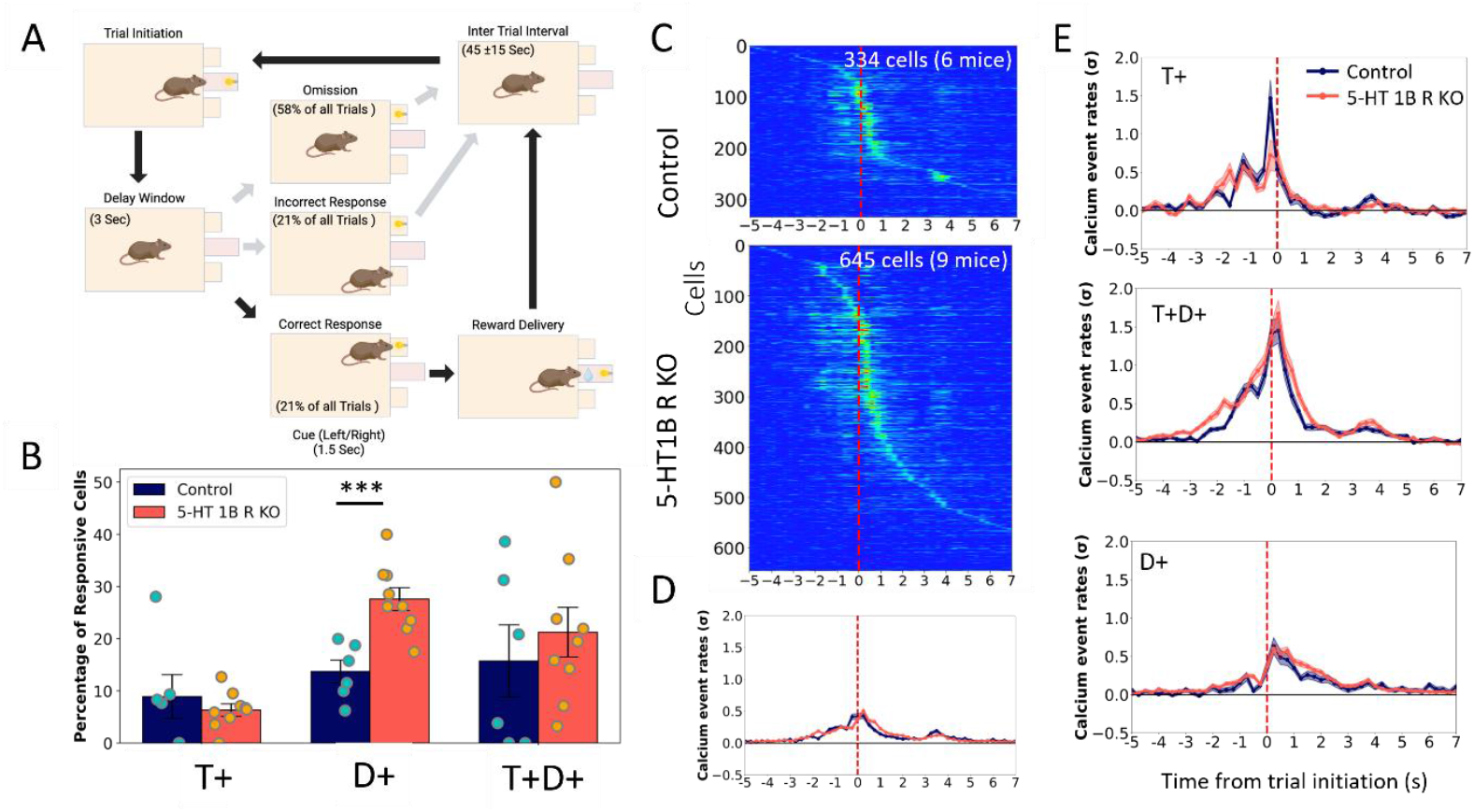
Mice lacking 5-HTlBRs have more active MSNs during the delay period. (A) Schematic showing the two-choice serial reaction time task including average percentage of trial types (omission, incorrect and correct responses) across all mice. (B) The percentage of responsive cells that show increases in calcium event rates during the trial initiation and/or delay window periods for both genotypes. (C) Normalized calcium event rates for cells averaged over all correct trials for Control (top) and 5-HT1BR KO mice (bottom), sorted by time of peak event rate. (D) Normalized event rates are shown averaged over all cells for Control (blue) and 5-HT1BR KO (orange) mice. (E) Traces of normalized event rates are shown for subsets of cells that were characterized as responsive to the trial initiation and/or delay period for both genotypes. *, p<0.001

We focused on cells that were responsive during the trial initiation and delay periods of the trial (Fig 5B), and found a significantly higher percentage of cells that were active during the delay period in mice lacking 5-HT1BR (t_13_=4.26, p=0.0009). There were no significant differences in the proportion of cells that increased their calcium event rates during the trial initiation period or through the trial initiation and delay period (t_13_=0.71, p=0.50 for T+ and t_13_=0.68, p=0.51 for T+D+). Looking at the calcium event rates over the trial for each of these categories of cells showed a similar pattern of increased calcium activity in mice lacking 5-HT1BR (Fig 5E). In the T+ cells, there was no effect of genotype on the average calcium event rates (F(1,54=0.018, p=0.894). However, for D+ and T+D+ cells, there were significant effects of genotype, with 5-HT1BR KOs showing higher calcium event rates during the trial (F_1,126_=11.58, p=0.0009 for T+D+; F1,124=4.68, p=0.033 for D+). Importantly, there were no significant effects of genotype on locomotion activity during the delay period (β=-259.9, SE=354.2, t_13_=-0.73, p=0.48), indicating that the increased calcium activity during the delay period did not likely stem from any differences in locomotor activity.

To further analyze the neural activity during the delay window, we trained mice for 6 sessions on this task to allow for improvement in performance. Both genotypes increased their performance on the task, showing increases in numbers of trials initiated (F_1,13_=29.41, p=0.0001) and rewarded (F_1, 13_=7.05. p=0.0198), though there was no change in the percentage of trials with premature responses (F_1,13_=0.37, p=0.555; Fig 6A). Focusing again on cells that showed increases in calcium event rates during the trial initiation and delay period, we find that there were no effects of session on the proportion of cells in each category (Fig 6B; F_1, 13_=0.9, p=0.359). Additionally, the genotype difference in the percentage of cells that are responsive during the delay window was diminished by the later sessions (F_1,13_=1.829, p=0.199). However, in both genotypes, the average calcium event rates of these cells increased significantly across the sessions (F_1, 217_=19.40, p<0.0001), with the event rates almost doubling from the first to the sixth session (Fig 6C). Comparing locomotor activity between the first and sixth sessions shows no significant difference in locomotor activity across sessions (β=-400.8, SE=312.2, t_28_=-1.28, p=0.21), matching the lack of change in premature responses), suggesting that the increased calcium activity during the delay period isn’t likely a consequence of any change in behavior during the task.

**Figure 6.**
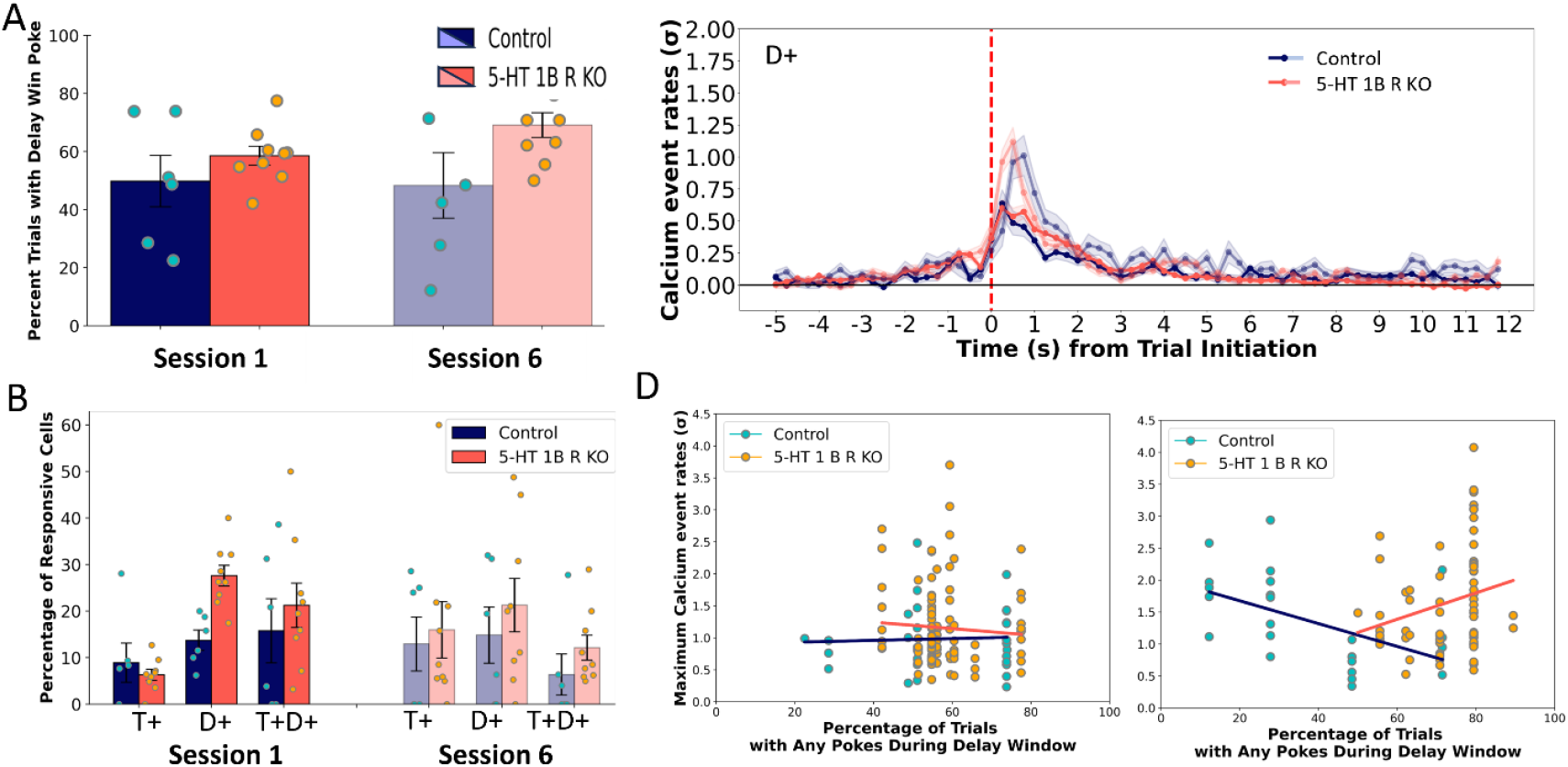
Training increases amplitude, but not proportion of delay window response neurons. (A) The percentage of responsive cells that show increases in calcium event rates during the trial initiation and/or delay window periods for both genotypes in the first and sixth training sessions of the waiting task. (B) Normalized calcium event rates for cells showing significant increases in calcium activity during the delay period, averaged over all correct trials for the first (darker) and sixth (lighter) sessions. (C) Behavioral performance is shown in terms of responses during the delay window for both genotypes of mice. (D) Scatterplots show correlations of the amplitude of the z normalized calcium event rate for individual cells showing increases in calcium activity during the delay period as a function of the number of impulsive responses made during the delay period for the first (left) and sixth (right) sessions. Lines of best fit are plotted for cells from mice from each genotype separately.

In order to examine the relationship between delay-period neural activity and behavior, we correlated the percentage of trials with premature responses with the maximum event rate of delay responsive cells for each animal (Fig 6D). During session 1 there was no correlation in either genotype (r=0.05; p=0.8 for controls; r=-0.07, p=0.49 for 5-HT1BR KOs) indicating that the calcium activity during the delay period was likely not purely a readout of response movement. However, during session 6, both genotypes showed significant correlations between behavior and calcium event rate, though interestingly in opposite directions. Controls showed a negative association with decreased event rates being associated with more premature responses (r=-0.51, p=0.012) while 5-HT1BR KOs showed increased event rates associated with more premature responding (r=0.24, p=0.048). These data suggest that over 6 sessions the calcium activity in the DMS changes differentially in the absence of 5-HT1BR expression, potentially underlying decreased inhibition of responding during the delay period.

## Discussion

Overall, our results show that 5-HT1BRs influence MSN activity in vivo during goal-directed behavior. While previous work has shown this effect in vitro using electrophysiology, our study was one of the first to examine the effects of 5-HT1BR during freely moving operant behavior using one-photon calcium imaging of MSNs in the DMS.

This allowed us to look at the influence of behavioral context on serotonin modulation of the DMS. Interestingly we found that the absence of 5-HT1BR expression resulted in differential effects depending on behavioral task requirements. The divergence is particularly remarkable given that the behavioral tasks were overall very similar—having the same general task structure, reward, and physical context. The difference was limited only to the waiting requirement during a delay period. First, in the task without a delay, we found that the absence of 5-HT1BR expression resulted in less excitatory calcium responses, and more inhibitory responses to reward in MSNs. This suggests that serotonin signaling through 5-HT1BRs can promote increased MSN responding to reward. On the other hand, in the second task which included a 3s delay period in the trial, we found that the absence of 5-HT1BR expression resulted in more cells showing excitation – suggesting that serotonin can inhibit MSN calcium activity via 5-HT1BRs during waiting. These nearly opposite effects of 5-HT1BR expression on MSN activity during responding and waiting were surprising given the overall similarity of the paradigm and trial structure.

Our results are consistent with past work implicating serotonin in regulating both approach and inhibition (Desrochers et al., 2022; Soubrié, 1986). A number of studies suggest that serotonin is necessary for patience. For example, depletion of serotonin increases premature responding, and optogenetic activation of serotonin neurons prolongs waiting time in rodents (K. Miyazaki et al., 2018; K. W. Miyazaki et al., 2014; Winstanley et al., 2004). Interestingly, serotonin is also important for behavioral vigor in the context of goal-directed responding (Bailey et al., 2018; Yoshida et al., 2019). In order to reconcile these two seemingly opposing ideas, one can consider that serotonin may act through different receptors, in different brain regions, or on different timescales in order to elicit the differential outcomes. However, in our studies, we imaged the same brain region in mice lacking a single serotonin receptor, and found opposite effects of 5-HT1BR signaling on DMS activity. One potential explanation that has been proposed previously (Roberts et al., 2020) is that serotonin acts differentially based on behavioral task context, therefore having varied effects depending on the state of the DMS influenced by other inputs. In the response activation state, serotonin signaling through 5-HT1BRs may promote DMS activity, while in the behavioral inhibition context, serotonin inhibits DMS excitability. In this way, serotonin acting through a single receptor in one brain region on similar timescales could contribute to promoting both approach and waiting behavior.

Given that we used mice lacking 5-HT1BR expression throughout the brain, our results do not speak to the location of the direct site of action of the effects of serotonin. It is possible that the receptors which influence MSN calcium activity are located on a number of cell types within the DMS, or even expressed on neurons at a distal brain region which send projections to MSNs (Burke & Alvarez, 2022; Cavaccini et al., 2018; Pommer et al., 2021; Virk et al., 2016). Furthermore, given that we used whole-life knockouts of 5-HT1BR, it is unclear when the effects of 5-HT1BR are occurring. There could be developmental effects resulting from of a lack of 5-HT1BR expression which are the proximate cause of the reported effects on calcium activity. Importantly, we chose to use the tetO1B mouse line in these studies (Nautiyal et al., 2015), so that future work can build upon this using inducible, cell-type-specific, tissue-specific, and projection-specific knockout approaches to investigate the mechanisms through which 5-HT1BRs influence MSN activity in the DMS during reward-related behaviors.

Another limitation of our study is that we did not image cell type-specific calcium activity of D1R- and D2R-expressing subtypes of MSNs in the DMS. We used the CaMKIIa promotor to drive expression of the calcium indicator in the DMS. Past characterizations of CaMKIIa expression has shown that it is expressed in the majority of MSNs and not expressed in ChAT, PV, NPY, or calretinin interneurons in the DMS (Klug et al., 2012). This suggests that the population of cells that we are imaging in the DMS includes both D1R- and D2R-expressing MSNs. Given mixed evidence supporting purported differential responses of direct and indirect pathway MSNs during goal-directed behavior, it is unclear if we would expect to see differences in effects between the two cell types. A number of papers show similar calcium responses of D1+ and D2+ cells encoding actions during goal-directed behavior, while others show evidence of the more classic opponent control these cell types in their encoding of action and reward (Bolkan et al., 2022; Cox & Witten, 2019; Cui et al., 2013; Kravitz et al., 2012; Malvaez et al., 2025; Shin et al., 2018; Tecuapetla et al., 2016).

Regardless of how D1+ and D2+ cells may encode goal-directed behavior differentially, it is still unknown if serotonin preferentially targets or differentially affects the activity of the cell types. However, there is limited evidence to support that signaling through 5-HT1BR expression would differentially influence the activity of D1+ versus D2+ MSNs. First, there is no evidence of differential expression of 5-HT1BRs on the different MSN subtypes (Pommer et al., 2021). Further, assuming that the effects seen are due to the absence of MSN-instrinsic 5-HT1BR expression in the DMS, we would not expect to see any differences on the effect of D1-vs D2-expressing MSNs given evidence that 5-HT1BR reduces lateral inhibition similarly in both cell-types (Burke & Alvarez, 2022; Pommer et al., 2021). Alternatively, if the effects of 5-HT1BR expression are via modulation of cortical inputs, we would again expect no cell-type specific effects based on evidence that 5-HT-LTD of MSNs is not pathway-specific (Mathur et al., 2011). Finally, we have previously shown that 5-HT1BR KO mice do not have altered release of dopamine in the DMS (Nautiyal et al., 2015), suggesting that 5-HT1BR expression could not differentially influence dopamine signaling on D1 vs D2 receptors in the DMS. Overall, future studies should directly test if there are differential effects of serotonin on direct and indirect pathway signaling.

In conclusion, our work addresses the effects of 5-HT1BRs on MSN activity in the DMS, in vivo. While many elegant electrophysiology studies have shown 5-HT1BR-dependent effects on MSN activity in vitro, examining these effects during behavior is important to understand how these effects are influenced by behavioral context. In particular, the functional role of serotonin signaling during decision making, goal-directed behavior, and other reward-related behavior is multi-faceted and likely modulated by setting. Additionally, given the multiple cell types and projections expressing 5-HT1BRs in the DMS, this is a likely site at which serotonin may differentially influence behavior dependent on the state of the DMS given that inputs are known to vary during approach and inhibition. Overall, our one photon calcium imaging data in the DMS show that 5-HT1BR modulation of MSN activity is dependent on behavioral state, and point to a need to study the effects of serotonin signaling on neural circuits in vivo to understand how context influences serotonin neuromodulation.

## Acknowledgments

Funding from R00 MH106731, R01 MH126178, and the William H. Neukom 1964 Institute for Computational Science CompX Faculty Award. We would like to thank Drs. Kyle Smith and Mitch Spring for their helpful comments

## Notes

### Competing Interest Statement

The authors have declared no competing interest.

